# DLL3 REGULATES NOTCH SIGNALING IN SCLC

**DOI:** 10.1101/2022.07.08.499386

**Authors:** Jun W. Kim, Julie H. Ko, Julien Sage

## Abstract

Tumor heterogeneity plays a critical role in tumor development and response to treatment. In small-cell lung cancer (SCLC), intratumoral heterogeneity is driven in part by the Notch signaling pathway, which reprograms neuroendocrine cancer cells to a less/non-neuroendocrine state. Here we investigated the atypical Notch ligand DLL3 as a biomarker of the neuroendocrine state and a regulator of cell-cell interactions in SCLC. We first built a mathematical model to predict the impact of DLL3 expression on SCLC cell populations. We next tested this model using a single-chain variable fragment (scFv) to track DLL3 expression *in vivo* and a new mouse model of SCLC with inducible expression of DLL3 in SCLC tumors. We found that high levels of DLL3 promote the expansion of a SCLC cell population with lower expression levels of both neuroendocrine and non-neuroendocrine markers. This work may influence how DLL3-targeting therapies are used in SCLC patients.

## INTRODUCTION

Intratumoral heterogeneity is a hallmark of cancer, with tumors consisting of a variety of cancer and non-cancer cells. This heterogeneity may vary over time and environmental conditions, including following therapy, and it can arise from genetic or epigenetic mechanisms. Intratumoral heterogeneity plays an important role in tumor development, including shaping the natural response of cancer cells to the immune system or treatment in the clinic (reviewed in (Black and McGranahan, 2021; Dagogo-Jack and Shaw, 2018; Jamal-Hanjani et al., 2015; Tammela and Sage, 2020)).

SCLC is an aggressive subtype of lung cancer with fast growth rates and a striking metastatic ability. The median overall survival of SCLC patients has remained close to 8-10 months in the past 3 decades, with a 5-year survival at ~6% and over 200,000 estimated deaths worldwide every year. Most tumors respond well initially to standard-of-care chemoradiation treatment, but resistance emerges rapidly in nearly all cases (reviewed in (Rudin et al., 2021)). The recent approval of T-cell immune checkpoint inhibitors has been beneficial to only a small fraction of SCLC patients (reviewed in (Barrows et al., 2022)). There is a critical need to develop biomarkers for SCLC development and more efficient therapeutic strategies.

Inter- and intratumoral heterogeneity are prominent in SCLC. SCLC tumors have been classified into different subtypes based on transcriptional programs driven by specific transcription factors, with the SCLC-A (expressing the ASCL1 transcription factor) and SCLC-N (expressing the NEUROD1 transcription factor) neuroendocrine subtypes being the most frequent in patients (Baine et al., 2020; Gay et al., 2021; Rudin et al., 2019). Importantly, mounting evidence indicates that individual SCLC tumors are comprised of heterogeneous populations of cancer cells with different levels of neuroendocrine gene programs (Calbo et al., 2011; Lim et al., 2017; Williamson et al., 2016). As an example of this intratumoral heterogeneity, in SCLC-A tumors, activation of Notch signaling can reprogram neuroendocrine (NE) cancer cells towards less/non-neuroendocrine (non-NE) phenotypes. The non-NE cells are less tumorigenic and genetic and pharmacologic approaches to promote NE to non-NE differentiation have been shown to decrease SCLC tumor growth (Augert et al., 2019). But non-NE can support the survival of the NE cells, including when tumors are treated with chemotherapy (Lim et al., 2017). In this context, the mechanisms activating Notch signaling are not completely understood, but may depend on the YAP transcriptional regulator (Shue et al., 2022). As another example, in SCLC-N tumors, expression of c-MYC can activate Notch signaling, which also promotes the transition of these tumors to a less neuroendocrine phenotype (Ireland et al., 2020). These data and data from human tumors indicate that both inter- and intratumoral heterogeneity are likely key contributors of the ability of SCLC tumors to become resistant to therapies (Gay et al., 2021; Stewart et al., 2020; Sutherland et al., 2022).

DLL3 (Delta-like ligand 3) is an atypical ligand for NOTCH receptors initially studied for its role in early pattern formation in mouse embryos (Bulman et al., 2000; Dunwoodie et al., 1997; Kusumi et al., 1998). Early work showed that DLL3 is an inhibitor of Notch signaling in a cell autonomous manner (Ladi et al., 2005), possibly in a cis-inhibition mechanism in the Golgi (Chapman et al., 2011; Geffers et al., 2007). However, genetic studies of DLL3 O-fucosylation indicate that interactions with NOTCH may not fully account for the physiological function of DLL3, suggesting possible Notch-independent roles for DLL3 *in vivo* (Serth et al., 2015). The mouse *Dll3* gene is a direct target of ASCL1 (Nelson et al., 2009) and *DLL3* is expressed in a significant number of human SCLC tumors. Importantly, Saunders and colleagues showed that DLL3 is present at the surface of SCLC cells and that targeting DLL3-expressing SCLC cells using an antibody-drug conjugate (ADC) can eradicate SCLC in pre-clinical models (Saunders et al., 2015). Clinical trials using Rovalpituzumab tesirine (Rova-T), an ADC targeting DLL3, have been unsuccessful largely due to toxic side effects of this molecule (e.g., (Blackhall et al., 2021; Johnson et al., 2021; Malhotra et al., 2021)). However, other strategies are being developed to target DLL3 expressing cells, (e.g., (Giffin et al., 2021; Hipp et al., 2020; Lakes et al., 2020; Tully et al., 2022)) and DLL3 remains a target of interest for detection and treatment of SCLC.

Here we developed tools and models to further investigate the role of DLL3 as a biomarker of SCLC and as a regulator of Notch signaling in SCLC. We used mathematical modeling based on published data to predict the potential role that DLL3 expression may have on tumor heterogeneity. We further developed a new single-chain fragment variable (scFv) that can bind DLL3 on cells in culture and *in vivo*. Finally, we used this scFv to test our mathematical models in a new genetically engineered mouse model of SCLC with inducible expression of DLL3. Our data show variable levels of DLL3 expression in SCLC and indicate that DLL3 expression contributes to intratumoral heterogeneity.

## RESULTS

### Modeling the role of DLL3 in Notch-driven intratumoral heterogeneity in SCLC

DLL3 is often viewed as a cis-inhibitor of Notch signaling during development because of its role in localizing NOTCH receptors in the Golgi (**Figure 1A**). To determine if DLL3 could suppress Notch activity in SCLC cells, we used cells derived from tumors in the *Rb^fl/fl^;p53^fl/fl^;Rbl2^fl/fl^;Hes1^GFP/+^* mouse model of SCLC (*RPR2;Hes1^GFP^* model, also known as *TKO;Hes1^GFP^*, for triple knockout) (Lim et al., 2017; Shue et al., 2022). In this model, SCLC cells express GFP when the *Hes1* promoter is active. Because *Hes1* is a target of Notch signaling, GFP expression can serve as a reporter of Notch activity, and HES1^GFP^-positive (HES1^pos^) cells are less/non-neuroendocrine (Lim et al., 2017; Shue et al., 2022). We isolated HES1^pos^ cells from *RPR2;Hes1^GFP^* mutant tumors, and ectopic expression of FLAG-tagged DLL3 in these HES1^pos^ cells led to fewer GFP-positive cells, providing further evidence that DLL3 can suppress Notch signaling in *cis* in this SCLC context (**Figure 1B-D** and **Supplementary Figure S1A**).

**Figure 1.**
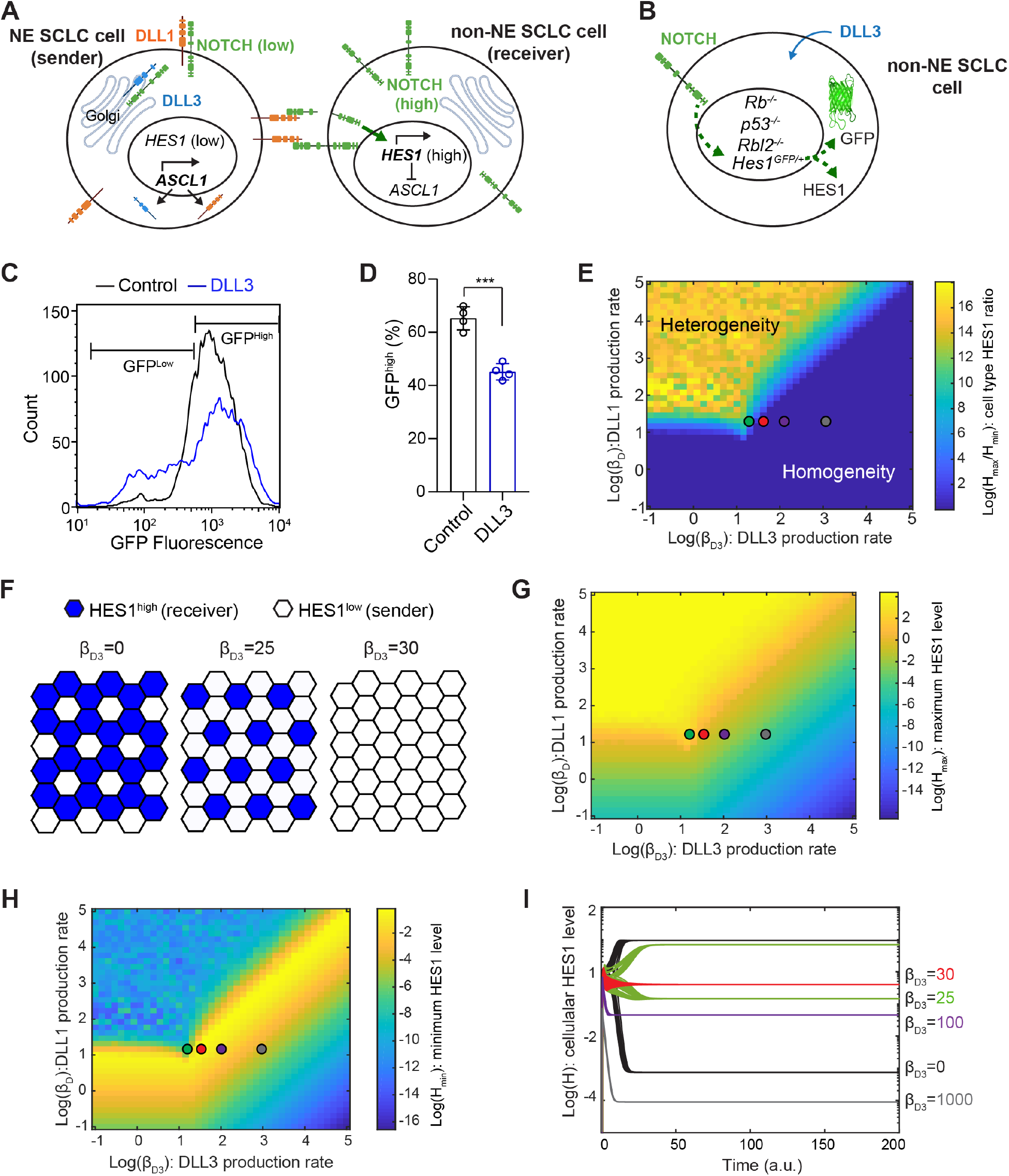
Mathematical model of mutual inactivation shows production rate dependent role of DLL3. (**A**) Schematic of Notch, DLL1, and DLL3 interaction in lateral inhibition with mutual inactivation (LIMI). (**B**) Schematic of HES1^GFP^-positive cells from an *Rb;p53;Rbl2;Hes1^GFP^* tumor serving as a reporter cell line, with GFP expression from the *Hes1* locus to monitor the effect of DLL3 expression in regulating Notch activity. (**C**) Flow cytometry of HES1^GFP^-positive cells with (blue) and without (black) ectopic expression of DLL3 (representative of n = 4 biological replicates). (**D**) Percentage of GFP^high^ HES1^GFP^-positive cells with and without ectopic expression of DLL3. Unpaired t-test, data represented as mean ±s.d. ***p<0.001 (**E**) Log(H_max_/H_min_) at steady state were calculated as a function of *β_D3_* and *β_D_.* Regions with values greater than 0 (light blue to yellow regions) support patterning, while those with 0 (dark blue) do not. (**F**) Hexagonal cell lattice with LIMI showing that DLL3 expression can lead to sparser patterns of higher HES1 cells (blue) or no pattern (homogeneous color). (**G**) H_max_ as a function of *β*_D3_ and *β*_D_. (**H**) H_min_ as a function of *β*_D3_ and *β*_D_. (**I**) Simulations with the indicated parameters for *β*_D3_ showing H levels in cells with high and low final H levels. Green, red, purple, and gray dots in (E), (G), and (H) correspond to the parameters used in (I).

In the Notch signaling pathway, activation of NOTCH receptors by their ligands, including Delta-like proteins (DLLs), leads to NOTCH cleavage, releasing the NOTCH intracellular domain (NICD). NICD induces transcription of the gene coding for HES1, which is an inhibitor of ASCL1, which is itself an activator of the expression of NOTCH ligands (DLLs, representing DLL1/3/4 and Jagged1/2 here) (Nelson et al., 2009; Ohsawa and Kageyama, 2008). This negative feedback leads to lateral inhibition and distinct cell fates among neighboring cells: low NOTCH/high DLL (‘sender’) cells and high Notch/low DLL (‘receiver’) cells (**Figure 1A**) (Collier et al., 1996). To gain a better understanding of the consequences of DLL3 expression on Notch signaling and intratumoral heterogeneity in SCLC, we built a simple mathematical model by incorporating DLL3 in the mutual inhibition model of lateral inhibition (Sprinzak et al., 2010) using the following reactions: (1) NOTCH receptors, denoted N, bind to DLL1 (as a representative of all NOTCH ligands, denoted D)in *trans,* which leads to cleavage of NOTCH and release of the Notch intracellular domain (NICD), denoted S; (2) NICD increases expression of HES1, denoted H (Bray 2006); (3) HES1 inhibits expression of DLL1 (Nelson et al., 2009; Ohsawa and Kageyama, 2008); (4) NOTCH receptors bind to DLL1 and DLL3, denoted D3, in *cis*, which inhibits NOTCH cleavage (Miller et al., 2009; Chapman et al., 2010; Ladi et al., 2005). As expected, setting the production rate of D3 at 0 replicated the results from previous works on mutual inactivation (Sprinzak et al., 2010; Sprinzak et al., 2011) supporting patterning even without cooperative regulatory feedback in the lateral inhibition model (**Supplementary Figure S1B-G**). We simulated this model using 1600 parameter sets across a two-dimensional parameter space spanning a wide range of production rates of D (*β_D_*) and D3 (*β_D3_*) (**Figure 1E**, **Equations 1–4** in Methods section, and **Table S1** for parameter values). To represent multicellular interactions, a two-dimensional hexagonal cell lattice was used for each parameter set. We plotted log(H_max_/H_min_), where H_max_ and H_min_ are the maximal and minimal HES1 levels in the hexagonal lattice at the steady state, respectively, as a function of *β_D_* and *β_D3_* to observe parameters leading to lateral inhibition, or heterogeneity in HES1 level (Formosa-Jordan and Sprinzak 2014).

The cis-inhibition model shows that when β_D3_ > β_D_, HES1 heterogeneity mostly does not occur, indicating effective inhibition of cell-type bifurcation (**Figure 1E**). Within the parameter regime leading to heterogeneity, β_D3_ also regulates the ratio of receiver (HES1^high^) cells and sender (HES1^low^) cells, with higher number of sender cells as β_D3_ increases (**Figure 1F**). We next examined H_max_ and H_min_ separately. H_max_ consistently decreases as β_D3_ increases (**Figure 1G**). Interestingly, a large region of parameter space shows higher H_min_ with increasing DLL3 (**Figure 1H,I**). H_min_ increases throughout the parameter regime that leads to heterogeneity, peaking near β_D3_ = β_D_, and gradually decreases as β_D3_ further increases (**Figure 1H**).

In this system, our mathematical modeling suggests that the influence of DLL3 on the heterogeneity of HES1 level depends on the level of DLL3 expression. At high levels (β_D3_ ≫ β_D_), DLL3 inhibits both heterogeneity and Notch activation. At lower levels, however, DLL3 allows trans-interaction between NOTCH and other DLLs while preventing lateral inhibition, which leads to homogeneous intermediate NOTCH activation throughout the cell population. In the parameter space leading to heterogeneity, DLL3 expression also affects sparsity of the receiver cells.

### Generation and validation of a single-chain fragment variable (scFv) binding DLL3

Given that differences in DLL3 expression level may contribute to intratumoral heterogeneity via NOTCH-driven patterning, we sought to examine whether DLL3 levels in tumors were related to the amount of NOTCH signaling or intratumoral heterogeneity present *in vivo.* To do this, we first developed a new tool to detect DLL3 in tumors *in vivo*.

Single-chain fragment variables (scFvs) have several advantages over antibodies, including their small size and their ease of production in bacteria (Peltomaa et al., 2022; Weisser and Hall, 2009). We generated a His-tagged scFv based on the sequence of an antibody targeting DLL3 (Methods) (**Figure 2A and Supplementary Figure S2A**). We first tested the binding of this scFv targeting DLL3 in 293T cells expressing exogenous DLL3 compared to controls. Flow cytometry analysis using an anti-His antibody showed increased binding in cells expressing mouse DLL3 (**Figure 2B,C**). Using the same approach, we were also able to detect endogenous DLL3 expression in human SCLC cell lines (**Figure 2C** and **Supplementary Figure S2B**). Thus, this scFv targeting DLL3 is capable of binding to DLL3 expressed on the surface of SCLC cells in culture.

**Figure 2.**
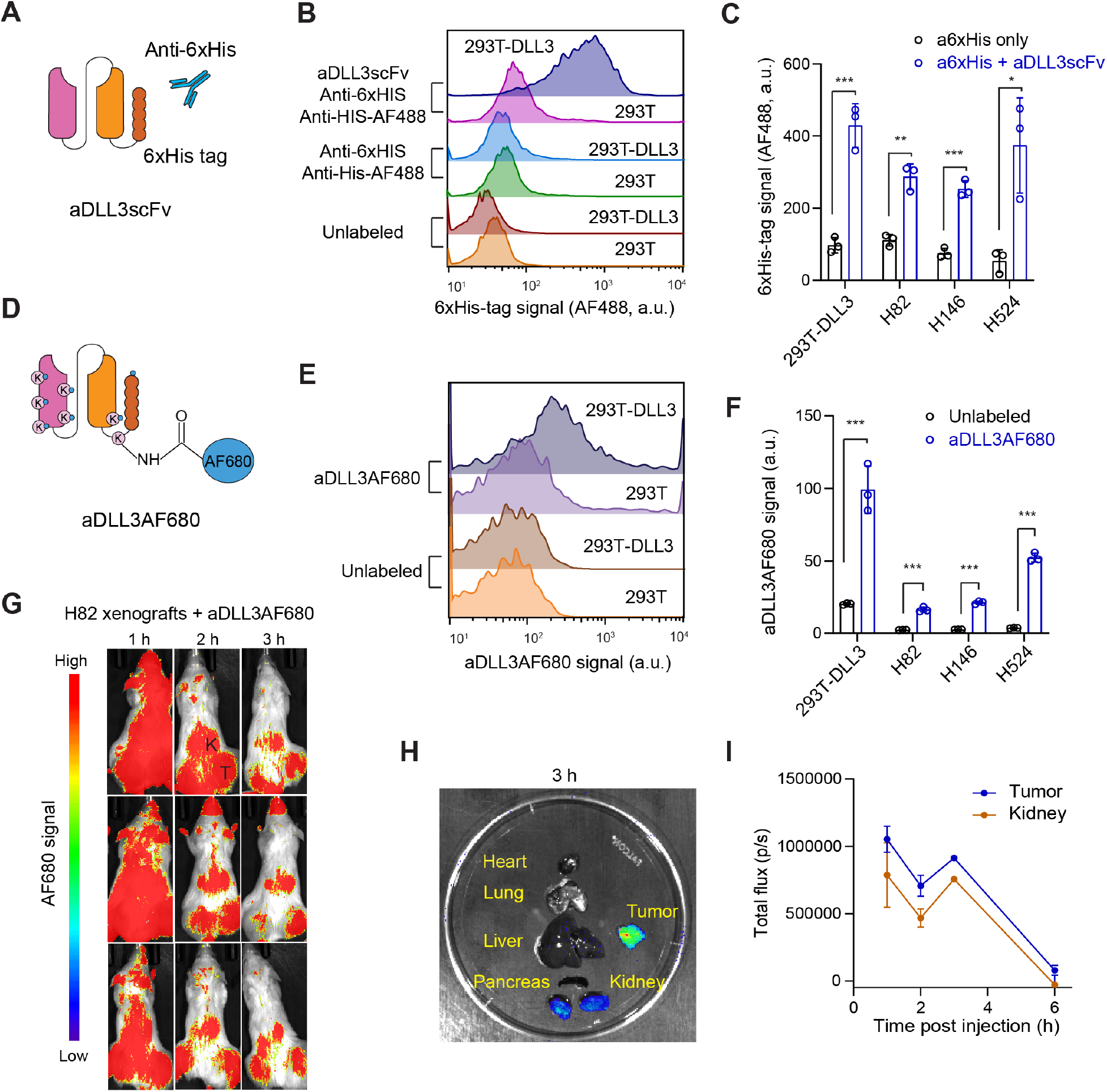
Detection of DLL3 expression at the surface of cells using anti-DLL3 scFv. (**A**) Schematic representation of the His-tagged scFv targeting DLL3 (aDLL3scFv) and the detection of cells expressing DLL3 using this molecule. (**B**) Representative flow cytometry analysis of DLL3 expression in 293T cells expressing exogenous mouse DLL3 using aDLL3scFv (from n=3 biological replicates). (**C**) Quantification of DLL3 detection in 293T-DLL3 cells from (B) and human SCLC cell lines. (**D**) Schematic representation of AF680 labeled aDLL3scFv (aDLL3AF680). (**E**) Representative flow cytometry showing binding activity of aDLL3AF680 (from n=3 biological replicates). (**F**) Quantification of (E) in human SCLC cell lines. (**G**) Whole body fluorescent images of H82 tumor xenografts with 2 nmol of AF680-labeled scFv injected via tail vein. Tumors (T) and kidneys (K) are indicated. (**H**) Representative *ex vivo* imaging of an NCI-H82 bearing mice in (H). (**I**) Quantification of imaging signals, reported as the total flux (p/s) (n=3). Unpaired t-test, data represented as mean ±s.d. p<0.05, **p<0.01, ***p<0.001.

To determine if this scFv molecule (aDLL3scFv) could be useful to detect SCLC tumors *in vivo,* we directly labeled the amino groups of lysine residues and the N-terminal group of the aDLL3scFv with Alexa Fluor 680 (AF680), a near-infrared fluorophore with strong tissue penetration that is suitable for *in vivo* imaging (**Figure 2D**). We first verified that the AF680-labeled aDLL3scFv (aDLL3AF680) still detected DLL3 on the surface of 293T cells expressing exogenous DLL3 as well as SCLC cells in culture (**Figure 2E,F** and **Supplementary Figure S2C**). Non-invasive optical imaging was performed over a 6-hour time period following tail-vein injections with 2 nmol of aDLL3AF680 in NSG mice bearing subcutaneous xenografts (NCI-H82 model). Whole-body fluorescent imaging showed prominent signal in the tumor area, as well as in the kidney (where the scFv molecules are excreted) 2-3 hours post injection (**Figure 2G,H**). Maximum tumor-to-normal tissue contrast was observed 2 hours post injection (**Supplementary Figure S2D**). Signal from the fluorescent scFv was strong after 1 hour and started to decrease after 3-4 hours, as expected for a small molecule with a short half-life (**Figure 2I**).

### Exogenous expression of DLL3 in mouse SCLC tumors

Notch signaling is a driver of non-neuroendocrine cell fates and intratumoral heterogeneity in SCLC (Ireland et al., 2020; Lim et al., 2017), and our mathematical modeling predicts that DLL3 may contribute to Notch signaling activity and may modulate heterogeneity. We next sought to test this idea experimentally in the *RPR2* mouse model of SCLC where this heterogeneity has been described (Lim et al., 2017; Shue et al., 2022). In this model, intra-tracheal instillation of an adenoviral vector expressing the Cre recombinase initiates tumors modeling the SCLC-A subtype of human SCLC. SCLC cells in these tumors express DLL3 at their surface (**Supplementary Figure S3A**).

To manipulate DLL3 levels in this model, we generated a new allele to conditionally induce DLL3 expression upon Cre-mediated recombination (*Rosa26^LSL-Dll3^* allele) (**Figure 3A**). We crossed this allele to the *RPR2* model *(RPR2;Rosa26^LSL-Dll3^* mice) and initiated tumors by Ad-CMV-Cre instillation. As controls, we used *RPR2;Rosa26^LSL-Luc^* mice (Jahchan et al., 2016). Tumors developed in *RPR2;Rosa26^Dll3^* mice, and their histology was indistinguishable from that of *RPR2;Rosa26^Luc^* mice; while we did not generate enough mice to rigorously quantify tumor number and tumor burden, we did not observe any striking difference in tumor development between the two models (**Figure 3B**). A cell line derived from a *RPR2;Rosa26^Dll3^* mutant lung tumor grew as spherical clusters in suspension typical of neuroendocrine SCLC cell lines (Lim et al., 2017) and showed weak but detectable expression of the tdTomato reporter linked to DLL3 expression, as well as the expected recombination of the *Lox-STOP-Lox* cassette, validating the new allele (**Figure 3A** and **Supplementary Figure S3B,C**).

**Figure 3.**
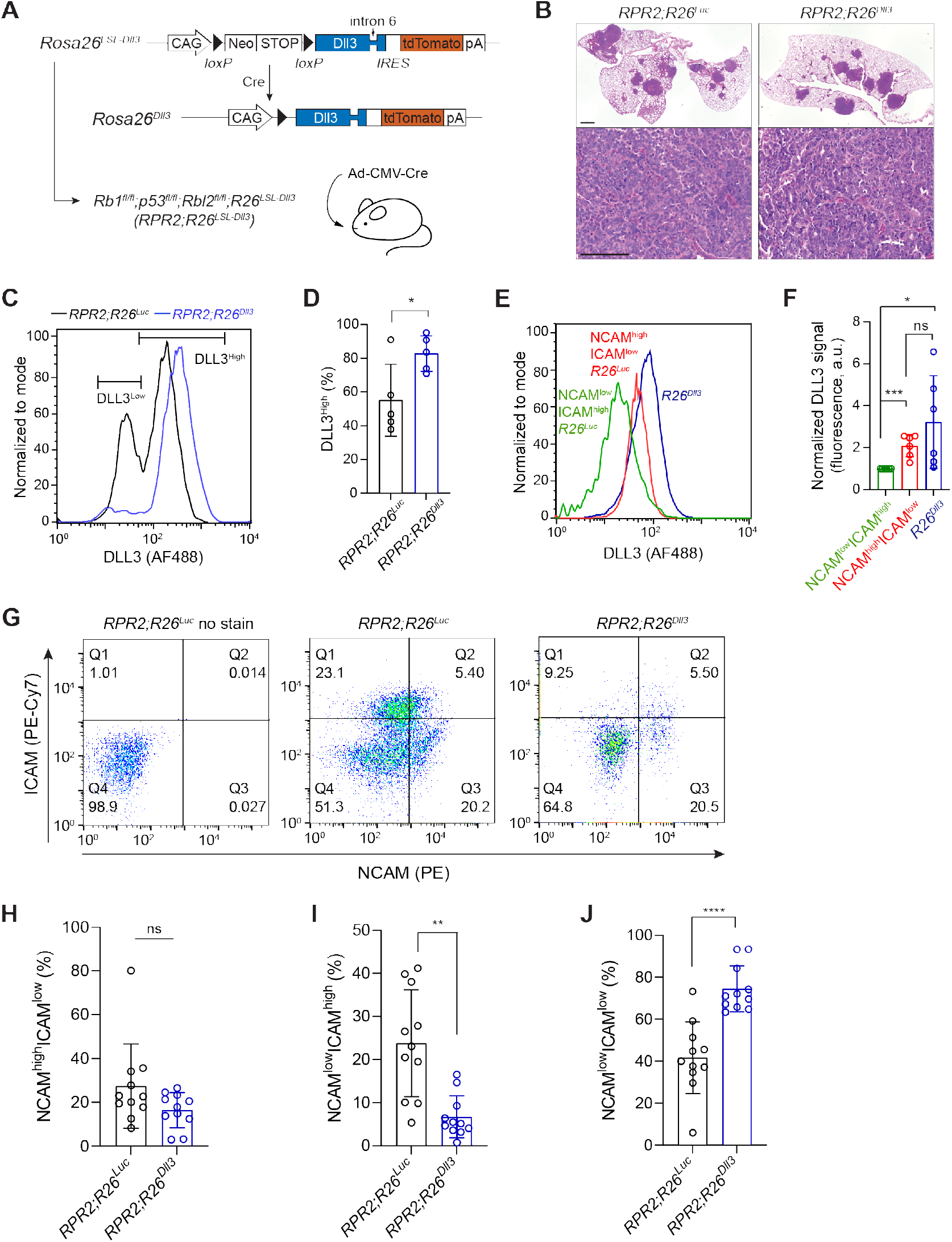
DLL3 expression perturbs the balance between neuroendocrine and non-neuroendocrine cells in a mouse model of SCLC. (**A**) Schematic representation of the Cre-inducible *Dll3* allele and the genetically engineered mouse model of SCLC. (**B**) Representative hematoxylin and eosin (H&E) staining of representative sections from *RPR2;R26^Luc^* and *RPR2;R26^Dll3^* mutant lungs (top, scale bar 1mm) and lung tumors (bottom, scale bar 100μm) 6 months after tumor initiation (n ≥ 6 mice). (**C**) Representative flow cytometry analysis of DLL3 cell surface expression in SCLC cells from *RPR2;R26^Luc^* and *RPR2;R26^D113^* mutant mice 5.5 months after tumor initiation. (**D**) Quantification of (**C**) (n=5 independent experiments). (**E**) As in (C) with *RPR2;R26^Luc^* SCLC cells differentiated by NCAM and ICAM expression. (**F**) Quantification of (E) (n=6 independent experiments). (**G**) Representative flow cytometry dot plots of control cancer cells from *RPR2;R26^Luc^* tumors with no stain, stained cells from *RPR2;R26^Luc^* tumors, and stained cells from *RPR2;R26^Dll3^* tumors (from left to right). (**H-J**) Quantification of NCAM^high^ ICAM^low^, NCAM^low^ ICAM^high^, and NCAM^low^ ICAM^low^ populations in (G). Unpaired t-test, data represented as mean ±s.d. p<0.05, **p<0.01, ***p<0.001.

We analyzed tumors from *RPR2;Rosa26^Dll3^* and *RPR2;Rosa26^Luc^* mutant mice ~5-6 months after tumor initiation for DLL3 expression using aDLL3scFv. In control *RPR2;Rosa26^Luc^* tumors, we observed two populations of cancer cells, as would be expected for tumors with neuroendocrine (high DLL3) and less/non-neuroendocrine (low DLL3) cells (**Figure 3C,D**). We validated this using NCAM and ICAM staining, with NCAM^high^ ICAM^low^ cells representing neuroendocrine cells and NCAM^low^ ICAM^high^ cells representing less/non-neuroendocrine, as validated previously by RNA-sequencing of these two populations from *RPR2* tumors (Shue et al., 2022) (**Figure 3E,F**). In *RPR2;Rosa26^Dll3^* mutant tumors, we observed fewer cells with low levels of DLL3 and a more homogeneous population of cells expressing intermediate/high levels of DLL3 (**Figure 3C,D**), indicating that the *Rosa26^Dll3^* allele can elevate DLL3 expression on the surface of DLL3^low^ cells but does not result in further overexpression on the surface of DLL3^high^ cells (**Figure 3C,D**). Similarly, while NCAM^high^ ICAM^low^ cells express higher level of DLL3 compared to NCAM^low^ ICAM^high^, no significant difference was detected between those of NCAM^high^ ICAM^low^ cells and cells from *RPR2;Rosa26^Dll3^* mutant tumors (**Figure E, F**).

We next quantified HES1 expression as a marker of intratumoral heterogeneity in *RPR2;Rosa26^Dll3^* and *RPR2;Rosa26^Luc^* mutant tumors, with HES1^high^ cells being Notch-active and less/non-neuroendocrine. As expected, we found regions with variable numbers of HES1-positive cells (Lim et al., 2017; Shue et al., 2022) (**Supplementary Figure S3D**). However, we did not detect any significant change in heterogeneity based on this marker between the two groups (**Supplementary Figure S3E**). While these experiments indicate that higher levels of DLL3 in tumors do not block the ability of SCLC cells to undergo a transition to a NOTCH-driven HES1-positive state, we reasoned that immunostaining may not be quantitative enough to identify more subtle changes.

We used flow cytometry to compare the expression of NCAM and ICAM in primary lung tumors from *RPR2;Rosa26^Dll3^* and *RPR2;Rosa26^Luc^* mice (**Figure 3G**). Compared to *RPR2;Rosa26^Luc^* tumors, we found that *RPR2;Rosa26^Dll3^* tumors exhibited a significant decrease in the NCAM^low^ ICAM^high^ non-neuroendocrine population with no significant change in the NCAM^high^ ICAM^low^ neuroendocrine population (**Figure 3G-I**). The change in the NCAM^low^ ICAM^high^ population was reflected by an increase in NCAM^low^ ICAM^low^ cells in *RPR2;Rosa26^Dll3^* tumors (**Figure 3G,J**).

The relatively homogeneous population in *RPR2;Rosa26^Dll3^* tumors suggests that higher expression of DLL3 in SCLC cells can inhibit the cell-fate bifurcation process mediated by the Notch signaling pathway. These observations support a model in which DLL3 contributes to Notch signaling activity in SCLC and may serve as a Notch pathway inhibitor in SCLC tumors *in vivo*.

## Discussion

Here we investigated the role of the atypical NOTCH ligand DLL3 in a mouse model of SCLC. In our three-pronged approach, we combined mathematical modeling, a new scFv molecule, and a new allele to induce DLL3 expression in mouse cells. We found that DLL3 expression contributes to the generation of specific subpopulations of SCLC cells, supporting a role for DLL3 in Notch signaling and in the control of intratumoral heterogeneity in SCLC.

While cell autonomous sources for intratumoral heterogeneity arise from various stressors and defects that generate genetic instability, non-cell autonomous contributors often involve signaling pathways that play critical roles during normal development and can modify the transcriptome of cancer cells. In this regard, a canonical process involves spatiotemporal regulation of the Notch signaling pathway. The Notch pathway is a highly conserved signaling mechanism implicated in both normal development and the progression of various cancer types. In different tissues, Notch signaling provides a binary fate switch through a biochemical feedback mechanism known as lateral inhibition, which regulates differentiation into different cancer cell types from an initially homogenous field of cells (reviewed in (Chitnis, 1995; Sjöqvist and Andersson, 2019)). Thus, the Notch signaling pathway promotes cell fate “bifurcation” rather than a specific cell fate. This contribution of Notch-mediated lateral inhibition to intratumoral heterogeneity has been investigated extensively in multiple settings, including breast cancer (Bocci et al., 2019), glioblastoma (Lim et al., 2015), and bladder cancer (Torab et al., 2021). To examine a potential role of DLL3 in SCLC in the classical Notch circuit, we incorporated DLL3 in the lateral inhibition model with a mutual inactivation model. Our mathematical modeling predicts that DLL3 expression can affect Notch signaling in several ways. As expected, when the production rate of DLL3 exceeds that of DLL1 (β_D3_ > β_D_) heterogeneity sharply decreases, indicating that DLL3 effectively reduces lateral inhibition (**Figure 1E**). Within the parameter regime of heterogeneity, DLL3 expression modulates the relative number of HES1^pos^ cells, making them sparser with increased DLL3 expression (**Figure 1F**). Interestingly, while very high production rates of DLL3 (β_D3_ ≫ β_D_) lead to inhibition of Notch activation in the entire population, over a wide range of parameters DLL3 expression maintains an intermediate level of Notch activation, which peaks near β_D3_ = β_D_ (**Figure 1H,I**). This observation reflects two seemingly opposing effects on Notch signaling. Low levels of DLL3 are sufficient to inhibit lateral inhibition, reducing or preventing the rise of HES1^low^ cells. At very high levels, DLL3 inhibits Notch activation entirely through complete cis-inhibition. In between these two states, the trans-activation of Notch by DLL1 reinforces intermediate Notch activity, even within the parameter regime that leads to the homogenous state. Thus, in the context of SCLC, our theoretical framework predicts that while high levels of DLL3 promotes the HES1^low^ phenotype (neuroendocrine), at lower levels it can lead to a hybrid phenotype that is neither HES1^low^ nor HES1^high^. When we examined the role of DLL3 experimentally in these processes, we found that ectopic expression of DLL3 in HES1^pos^ cells results in decreased HES1 expression, supportive of a cis-inhibitory function for DLL3. Intriguingly, induction of DLL3 in a mouse model of SCLC revealed that this ectopic expression decreases the relative number of cells in the non-neuroendocrine (NCAM^low^ ICAM^high^) population and increases that of a NCAM^low^ ICAM^low^ population, without significantly altering the neuroendocrine (NCAM^high^ ICAM^low^) population of cancer cells. This result suggests that DLL3 inhibits the bifurcation of SCLC cell fates. In future studies, it will be interesting to examine the exact nature of the NCAM^low^ ICAM^low^ population in *RPR2;Rosa26^Dll3^* tumors, such as whether these cells represent a precursor cell similar to lung epithelial cells during early lung development before Notch signaling is activated or another population of non-NE cells that do not require Notch (Hong et al., 2022). Our new mouse allele for DLL3 expression may also be useful to investigate how DLL3 plays Notch-dependent and Notchin-dependent roles both in developmental processes and in cancer.

Because several therapeutic approaches currently in clinical development rely on the expression of DLL3 on SCLC cells, validating its expression before and during treatment could benefit patients with SCLC. Monitoring DLL3 levels in SCLC cells could be used to track overall progression of SCLC and, as our data suggest, heterogeneity between SCLC cell subpopulations. PET imaging of DLL3 has been evaluated clinically using DLL3 targeting antibodies with promising results (Sharma et al., 2017). However, slow normal tissue clearance and requirement of radioisotopes with long half-lives hamper clinical application of antibodies as noninvasive imaging agents (Wei et al., 2020). Here, we generated an anti-DLL3 scFv (~1/8^th^ the size of an antibody) labeled with a fluorophore (aDLL3AF680) with tissue penetration, allowing noninvasive optical imaging. aDLL3AF680 successfully labeled SCLC tumors *in vivo* with good tumor-to-normal tissue contrast in a short time frame (2 hours post injection). Fast clearance may lead to a superior imaging agent for clinical application. Small size may allow better tissue penetration. Thus, scFv DLL3 binder may be a great alternative to full-length antibodies for *in vivo* detection and targeting of DLL3 in SCLC and other tumor types expressing this molecule on their surface.

## Methods

### Ethics statement

Mice were maintained according to practices prescribed by the National Institute of Health at Stanford’s Research Animal Facility (APLAC protocol #13565) and by the Institutional Animal Care and Use Committee (IACUC) at Stanford. Additional accreditation of Stanford Research Animal Facility was provided by the Association for Assessment and Accreditation of Laboratory Animal Care.

### Synthesis of anti-DLL3 scFv

*Escherichia coli* BL21 (DE3) cells were transformed with anti-DLL3 scFv (Raum et al., 2017) with c-terminal hexahistidine tag in pET vector by VectorBuilder. The transformed cells were grown overnight in LB agar plates with 100 μg/mL ampicillin. A single colony from each plate was inoculated in LB broth containing ampicillin and grown overnight. The primary culture was diluted 1:100 and grown at 37°C. When OD_600_ reached 0.5, the cells were induced with IPTG 0.5 mM at 37°C for 6 h. The cells were then harvested by centrifugation at 3,500 g, and the cell pellet was re-suspended with B-PER reagent (Thermo Fisher Scientific 90084) with lysozyme (0.1 mg/mL) and DNase I (50 μg/mL) and incubated for 30 min at 37°C. Lysates were centrifuged at 15,000 × *g* for 10 min, and the supernatants were purified using a nickel-NTA affinity column (Qiagen 30210). Centrifugal filters with molecular 10 kDa molecular weight cutoffs (Millipore UFC801024) were used for further purification and buffer exchange into PBS. Protein purity was further analyzed using sodium dodecyl sulfate polyacrylamide gel electrophoresis and quantified using a plate reader.

### AF680 dye conjugation

Purified scFv was buffer-exchanged to 0.1 M sodium bicarbonate buffer (pH 8.3) and concentrated to 3 mg/mL. AF680 succinimidyl ester (Thermo Fisher Scientific A20008) was dissolved in DMSO at 10 mg/mL and added to the protein solution to give a final concentration of 1 mg/mL. The reaction was incubated for 1 hr at room temperature. Unreacted dye removal and buffer exchange to PBS was done using a 10 kDa centrifugal filter.

### Cell binding assays

5×10^5^ human and mouse cells were incubated with 50 μL of 30 μg/mL aDLL3AF680 for 1 hr in PBS with 0.1% BSA (PBSB) at 4°C. The cells were then washed twice with PBSB and analyzed by flow cytometry. For testing unlabeled aDll3scFv, cells were incubated with 50 μL of 30 μg/mL aDll3scFv for 1 hr in PBSB at 4°C. The cells were washed once with PBSB, and secondary binding was performed on ice for 30 min using rabbit anti-hexahistidine antibody (Cell Signaling Technology 12698S) diluted 1:100 in PBSB. The cells were washed once with PBSB and incubated with goat anti-rabbit antibody labeled with Alexa Fluor 488 (Invitrogen A11008) diluted 1:100 in PBSB for 30 min on ice. The cells were then washed twice with PBSB before being analyzed by flow cytometry.

### Cell culture

Cell lines were cultured in RPMI-1640 (Corning MT15040CV) except for 293T cells and cell lines derived from *RPR2;R26^Luc^* and *RPR2;R26^Dll3^,* which were cultured in DMEM (Hyclone SH30243.01). RPMI and DMEM were supplemented with 10% bovine growth serum (Hyclone SH3054103HI) and penicillin-streptomycin-glutamine (Gibco 10378016). Cells were grown at 37°C in standard cell culture incubators. All cells were routinely tested (MycoAlert Detection Kit, Lonza) and confirmed to be free of mycoplasma contamination.

NCI-H524, NCI-H82, and NCI-H1694 cells were purchased from ATCC. Murine KP11B6 (a negative control for tdTomato expression in **Supplementary Figure S3B**) and HES1^GFP/+^ SCLC cell lines were generated in the lab and have been described (Lim et al., 2017).

### Lentiviral transduction

5×10^5^ 293T cells were seeded in each well of a 6-well plate and left to adhere overnight in antibiotics-free DMEM High-Glucose medium supplemented with 10% bovine growth serum. Lentiviral vectors were co-transfected with delta8.2 and VSV-G lentiviral packaging vectors using PEI. The supernatant was collected after 48 hr and added to the target cells. Cells were treated on day 5 with the selection agent according to the lentiviral vector used.

### Plasmids/sequences

For mammalian expression of DLL3, mouse DLL3 with a C-terminal FLAG-tag was cloned into the pCDH lentiviral expression vector.

### Flow cytometry

Flow cytometry was performed using a 100 μm nozzle on BD FACSAria II (Stanford Stem Cell Institute FACS Core) and analyzed with the FACSDiva software. For single cell analysis of lung tumors, a sequential gating strategy was used to analyze cancer cells by staining with the following FACS antibodies (**Figure S3F**): CD45-Pacific Blue (BioLegend 103126, 1:100), CD31-Pacific Blue (BioLegend 102422, 1:100), TER-119-Pacific Blue (BioLegend 116232, 1:100), CD24-APC (eBioscience 17-0242-82, 1:200), ICAM1-PE-CY7 (BioLegend 116122, 1:100), and NCAM-PE (R&D Systems FAB7820P, 1:100) (Shue et al., 2022). Data were analyzed using Flowjo software.

### Lateral inhibition with mutual inactivation (LIMI) model with DLL3

Notch receptor (N_i_) in cell *i* interacts with DLL1 (D_j_) on the surface of a neighboring cell *j*. This *trans*-activation leads to cleavage of the Notch receptor, which frees the intracellular domain, NICD (S_*i*_) to induce expression of downstream genes such as Hes1 (H_i_), described by the class lateral inhibition model (Collier et al., 1996). Notch receptor (N_i_) also interact with DLL1 (D_i_) on the same cell surface, which leads to cis-inhibition, described by the LIMI model (Sprinzak et al., 2010). In order to see the effect of cis-inhibition by DLL3 (D3_i_), Notch receptor (N_i_), and DLL1 (D_i_) on the same cell surface, we modified LIMI by considering the following system in two-dimensional array:

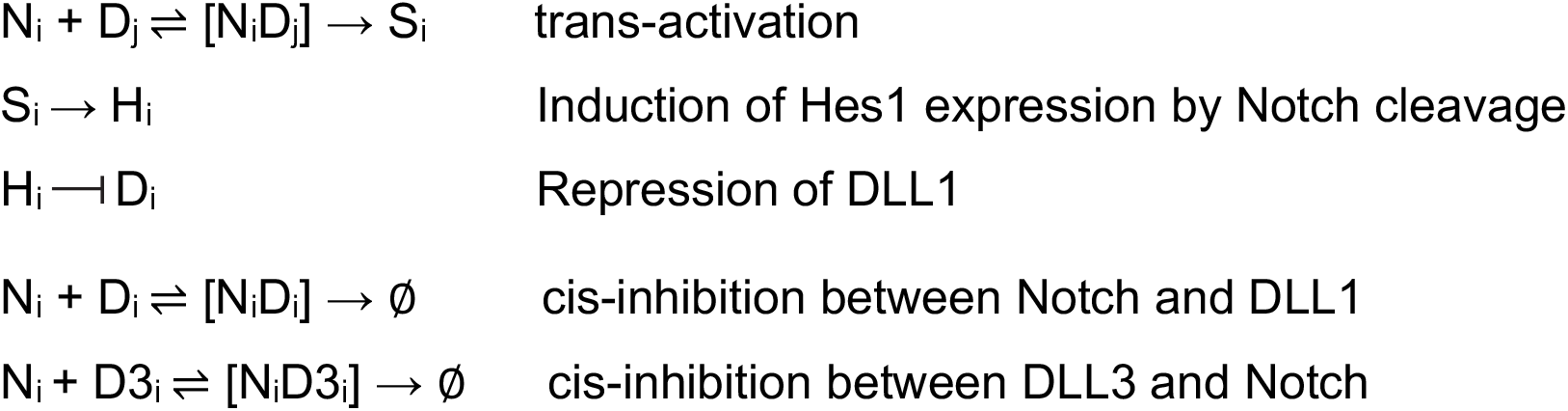

The first four reactions that do not describe DLL3 are directly from LIMI (Sprinzak et al., 2010). This model can be described by the following set of equations applied to a two-dimensional hexagonal lattice of cells:

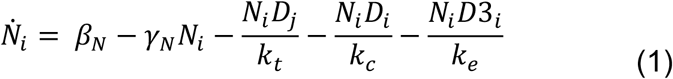

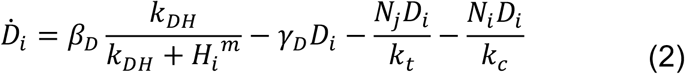

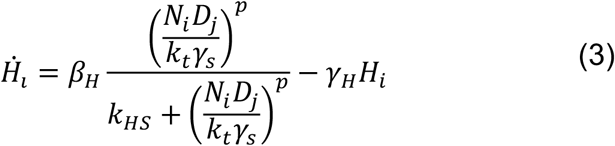

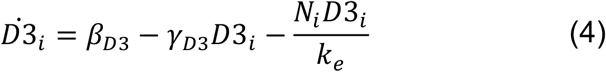

Equations 1–3 are from LIMI (Sprinzak et al., 2011). N_i_ and D_j_ interact at a rate k_t_^-1^. N_i_ and D_i_ interact at a rate k_c_^-1^. N_i_, D_i_, and H_i_ are produced at a rate of β_N_, β_D_, and β_H_, respectively. N_i_, D_i_, and H_i_ are degraded at a rate of γ_N_, γ_D_, and γ_H_, respectively. Repression of D_i_ by H_i_ is represented with a decreasing Hill function as a function of k_DH_ and m. Expression of H_i_ by S_i_ is represented with an increasing Hill function as a function of k_HS_ and p. Equation 4 describes D3_i_, where D3_i_ is produced, degraded, and interact with N_i_ at a rate β_D3_, γ_D3_, and k_e_, respectively. These equations are used in Figure 1 and Supplementary Figure 1. For Supplementary Figure S1B, D, and F, the mutual inactivation represented by 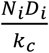 was omitted.

Numerical computations were performed using MATLAB’s ode15s solver (R2017b, MathWorks) using the scripts provided by Formosa-Jordan and Sprinzak (Formosa-Jordan and Sprinzak, 2014). The modified MATLAB codes can be downloaded from https://github.com/junwkim1/DLL3_lateral_inhibition_2022.

### Mouse models

The *Rb1^flox/flox^;Trp53^flox/flox^;Rbl2^flox/flox^ (RPR2,* or *TKO)* mouse model and the *Rosa26^lox-stop-lox-luciferase^* (*R26^Luc^*) allele have been previously described (Jahchan et al., 2016; Schaffer et al., 2010). Mice were maintained in a mixed C57BL/6;129SVJ background.

Cre-inducible DLL3-overexpressing mice (*R26^Dll3^*) were generated by knocking in a *Lox-Stop-Lox-Dll3-IRES-tdTomato* cassette into the *Rosa26* allele. The mice were generated by genOway via homologous recombination into the *Rosa26* allele in mouse embryonic stem cells, and clones were then injected into blastocysts to generate chimeric mice with germline transmission. The *Dll3* sequence in this allele is a cDNA/gDNA combo that retains intron 6 within the cDNA sequence to allow for expression of potential splice variants.

*R26^LSL-Dll3^* mice were crossed to *RPR2;R26^LSL-Luc^* mice to generate *RPR2;R26^LSL-Dll3/LSL-Luc^* mice, which were then crossed to each other to generate *RPR2;R26^LSL-Dll3/LSL-Dl13^* mice and *RPR2;R26^LSL-Luc/LSL-Luc^* littermate controls. Tumors were initiated at 8-12 weeks of age with intratracheal instillation of 4×10^7^ plaque-forming units of Adeno-CMV-Cre (Baylor College of Medicine, Houston, TX). Animals were euthanized and tumors were collected at ~6.5 months post-initiation, or earlier if they showed signs of respiratory distress. Mice were housed at 22°C ambient temperature with 40% humidity and a light/dark cycle of 12 hours (7am – 7pm).

### Genotyping

Mice were genotyped by using the Mouse Direct PCR kit (Bimake) on DNA isolated from tails following the manufacturer’s protocol.

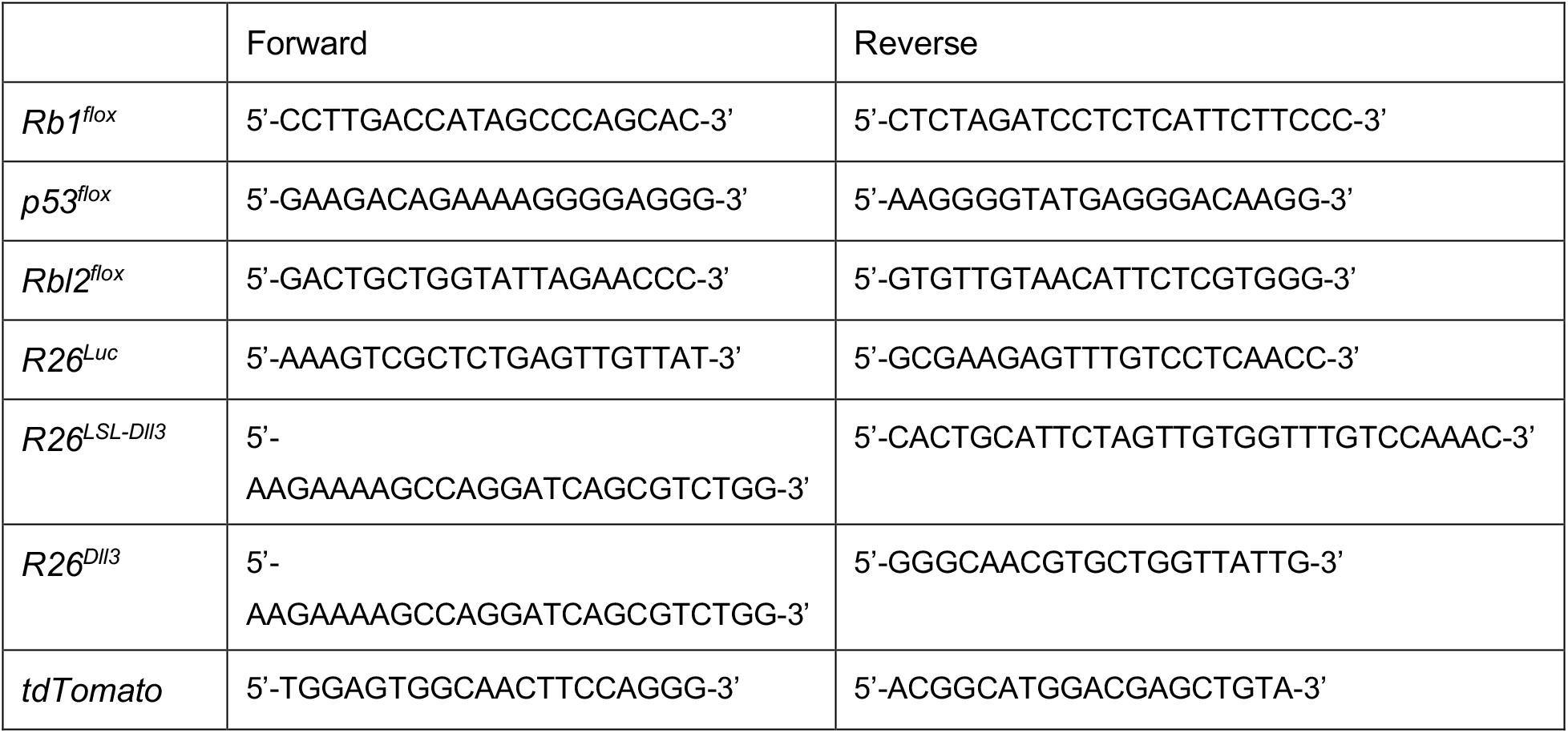

### Immunostaining

Lung lobes were fixed overnight in 10% formalin in PBS before paraffin embedding. Paraffin sections were deparaffinized with Histo-Clear (National Diagnostics HS-200) and gradually rehydrated from ethanol to water. Antigen retrieval was carried out in citrate-based unmasking solution (Vector Laboratories H-3300) by microwaving at full power until boiling, then 30% power for 25 min (DLL3) or 15 min (HES1), then left to cool at room temperature for 10 min before washing with water. Endogenous peroxidase was blocked by incubating slides in 3% hydrogen peroxide for 1 h. Sections were washed in PBST (PBS with 0.1% Tween-20), blocked in 5% horse serum for 1 h, and incubated with anti-DLL3 (1:200 Invitrogen PA5-23448) or anti-HES1 (1:200 Cell Signaling Technology 11988) diluted in PBST overnight at 4°C. DLL3 was developed using ImmPRESS^®^HRP Horse Anti-Rabbit IgG Polymer Detection Kit (Vector Laboratories MP-7401) following the manufacturer’s protocol. HES1 was developed using ImmPRESS^®^ Excel Amplified Polymer Staining Anti-Rabbit IgG Peroxidase Kit (Vector Laboratories MP-7601) following the manufacturer’s protocol. All sections used DAB substrate kit (Vector Laboratories SK-4100) for color development. Sections were counterstained with hematoxylin (Sigma-Aldrich HHS32-1L), gradually dehydrated from water to ethanol to xylene, and mounted with Refrax mounting medium (Anatech Ltd711). Sections were imaged using Keyence BZ-X700 all-in-one fluorescence microscope with BZ-X Viewer program version 1.3.1.1 and BZ-X Analyzer 1.4.0.1.

### Single-cell suspension

Tumors were dissected from lungs between 5 to 6.5 months after tumor induction, finely chopped with a razor blade, and digested for 15 min at 37°C in 10 mL of L-15 medium (Sigma L1518) containing 4.25 mg/mL Collagenase I (Sigma C0130), 1.4 mg/mL Collagenase II (Sigma 6885), 4.25 mg/mL Collagenase IV (Sigma 5138), 0.625 mg/mL Elastase (Worthington LS002292), and 0.625 mg/mL DNase I (Roche 10104159). The digested mixture was filtered through a 40 μm filter and centrifuged at 400 g for 5 min at room temp, and the resulting pellet was resuspended in 1 mL RBC lysis buffer (eBioscience 00-4333-57) for 30 sec, diluted in 30 mL PBS, and centrifuged at 400 g for 5 min at room temp. The pelleted cells were then resuspended in 0.1% BSA/PBS for flow cytometry or in cell culture media to generate cell lines.

### Statistical analysis

Statistical significance was assayed with GraphPad Prism software. Data are represented as mean ± sd. *p<0.05, **p<0.01, ***p<0.001, ****p<0.0001, ns not significant. The tests used are indicated in the figure legend.

## Data availability

All data are available in the article or from the corresponding author upon request.

## Acknowledgements

The authors thank Dr. Laura Saunders and Dr. Chen-Hua Chuang at Abbvie/StemCentrx for their help during the course of this study, including with the generation of the *R26^LSL-Dll3^* mice. We would like to thank David S. Glass for discussions and advice, as well as Myung Chang Lee for his comments on the manuscript. We also thank Pauline Chu from the Stanford Histology Service Center and all the members of the Sage lab for their help throughout this study. Research reported in this publication was supported by the Ludwig Institute for Cancer Research (J.S.), the NIH (grants CA217450, CA213273, and CA231997 to J.S., CA257169 to J.H.K.), and the A.P. Giannini Foundation (J.W.K.).

## Contributions

J.W.K., J.H.K, and J.S. designed the experiments, interpreted the results, and wrote the manuscript; J.W.K. performed the mathematical modeling and developed the scFv molecules; J.H.K performed experiments with mice; J.W.K. and J.H.K. worked together on experiments with cells in culture and tumor analysis.

## Competing interests

J.S. received research funding from StemCentrx/Abbvie for the development of the new *Dll3* allele. J.S. has equity in, and is an advisor for, DISCO Pharmaceuticals. The authors declare no other competing interests.

## Figures and Figure Legends

**Figure S1 related to Figure 1.**
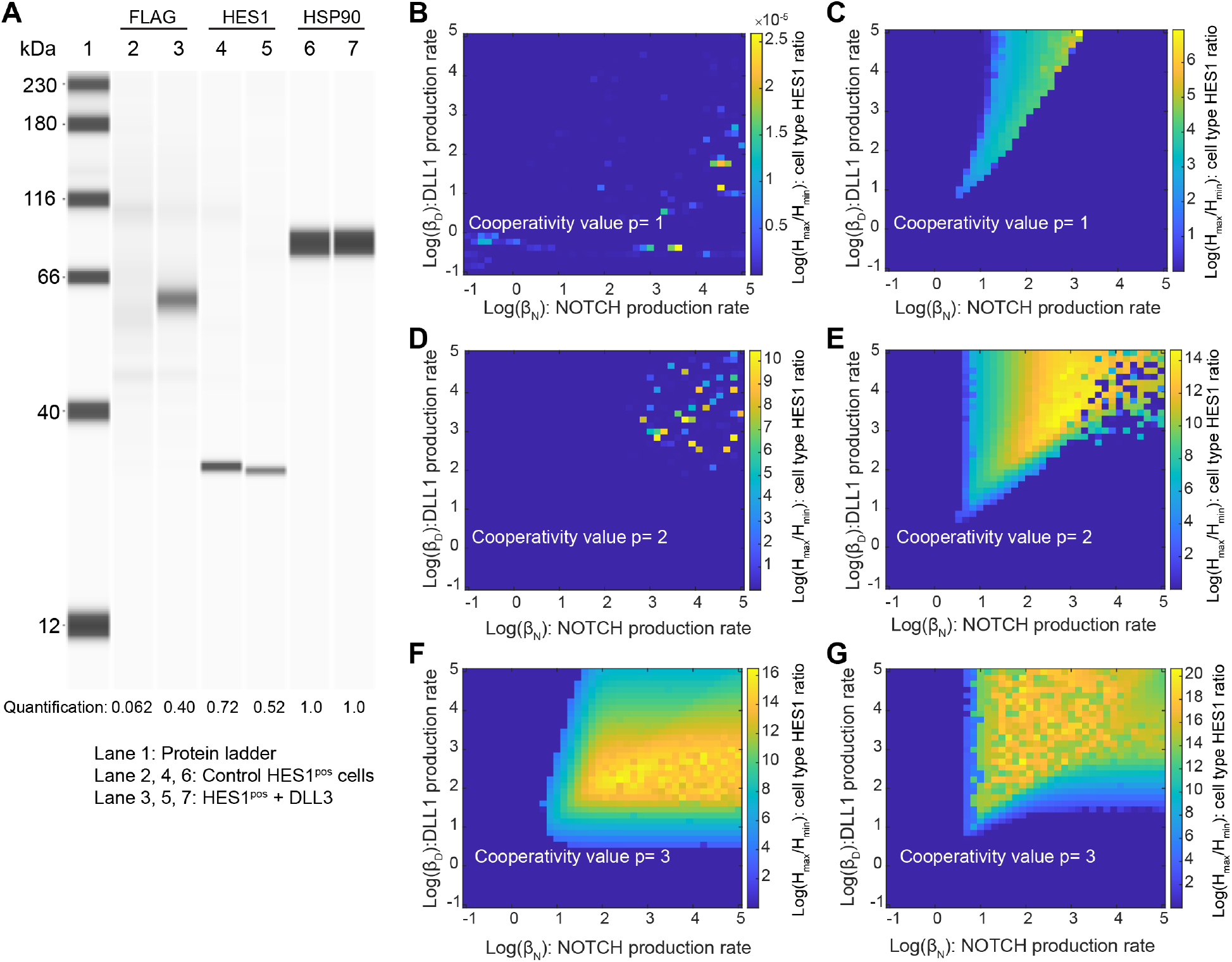
Ectopic expression of DLL3 in HES1^GFP^-positive cells. (**A**) Immunoassay measuring DLL3 and HES1 expression in control HES1^GFP^-positive cells and the same cells with stable expression of DLL3. HSP90 serves as a loading control. The expected molecular weights of DLL3, HES1, and HSP90 are 65 kDa, 30 kDa, and 90 kDa, respectively. The band intensities were quantified and normalized by those of HSP90. (**B-G**) Log(H_max_/H_min_) at steady state were calculated as a function of *β*_N_ and *β*_D_ without (**B, D, F**) and with (**C, E, G**) mutual inactivation using different values of the feedback loop cooperativity, p. Regions with values greater than 0 (light blue to yellow regions) support patterning, while those with 0 (dark blue) do not. As was shown in the studies by Sprinzak *et al.,* when p = 1 mutual inactivation was required to support patterning in the tested parameter space. Note the different scale in (**B**) (x10^-5^).

**Table S1 related to Figure 1:**
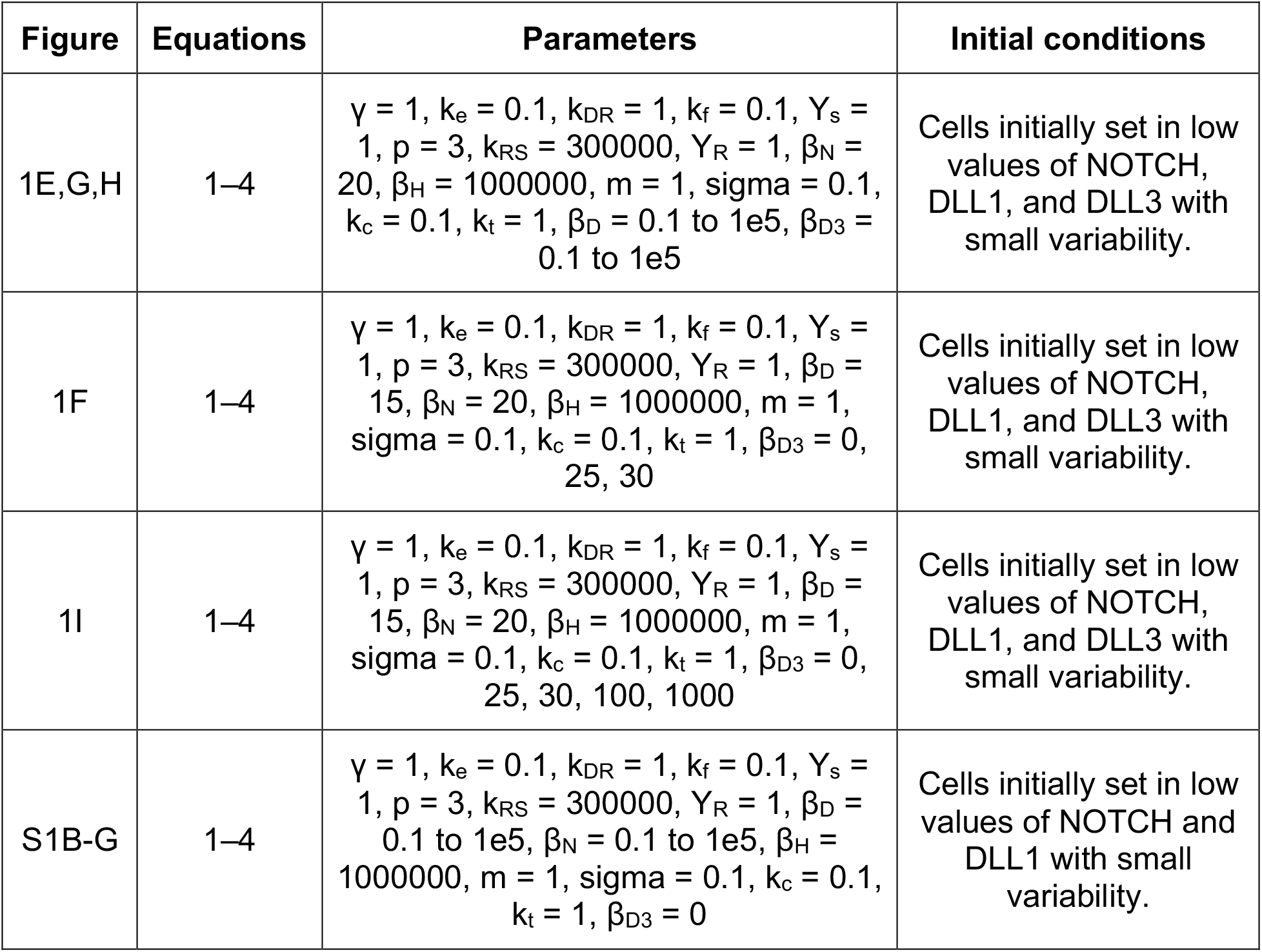
Parameter values for mathematical modeling. See equations in Methods and Figure 1

| Figure | Equations | Parameters | Initial conditions |
| --- | --- | --- | --- |
| 1E,G,H | 1–4 | $\gamma = 1, k_e = 0.1, k_{DR} = 1, k_f = 0.1, Y_s = 1, p = 3, k_{RS} = 300000, Y_R = 1, \beta_N = 20, \beta_H = 1000000, m = 1, \sigma = 0.1, k_c = 0.1, k_t = 1, \beta_D = 0.1 \text{ to } 1e5, \beta_{D3} = 0.1 \text{ to } 1e5$ | Cells initially set in low values of NOTCH, DLL1, and DLL3 with small variability. |
| 1F | 1–4 | $\gamma = 1, k_e = 0.1, k_{DR} = 1, k_f = 0.1, Y_s = 1, p = 3, k_{RS} = 300000, Y_R = 1, \beta_D = 15, \beta_N = 20, \beta_H = 1000000, m = 1, \sigma = 0.1, k_c = 0.1, k_t = 1, \beta_{D3} = 0, 25, 30$ | Cells initially set in low values of NOTCH, DLL1, and DLL3 with small variability. |
| 1I | 1–4 | $\gamma = 1, k_e = 0.1, k_{DR} = 1, k_f = 0.1, Y_s = 1, p = 3, k_{RS} = 300000, Y_R = 1, \beta_D = 15, \beta_N = 20, \beta_H = 1000000, m = 1, \sigma = 0.1, k_c = 0.1, k_t = 1, \beta_{D3} = 0, 25, 30, 100, 1000$ | Cells initially set in low values of NOTCH, DLL1, and DLL3 with small variability. |
| S1B-G | 1–4 | $\gamma = 1, k_e = 0.1, k_{DR} = 1, k_f = 0.1, Y_s = 1, p = 3, k_{RS} = 300000, Y_R = 1, \beta_D = 0.1 \text{ to } 1e5, \beta_N = 0.1 \text{ to } 1e5, \beta_H = 1000000, m = 1, \sigma = 0.1, k_c = 0.1, k_t = 1, \beta_{D3} = 0$ | Cells initially set in low values of NOTCH and DLL1 with small variability. |

**Figure S2 related to Figure 2.**
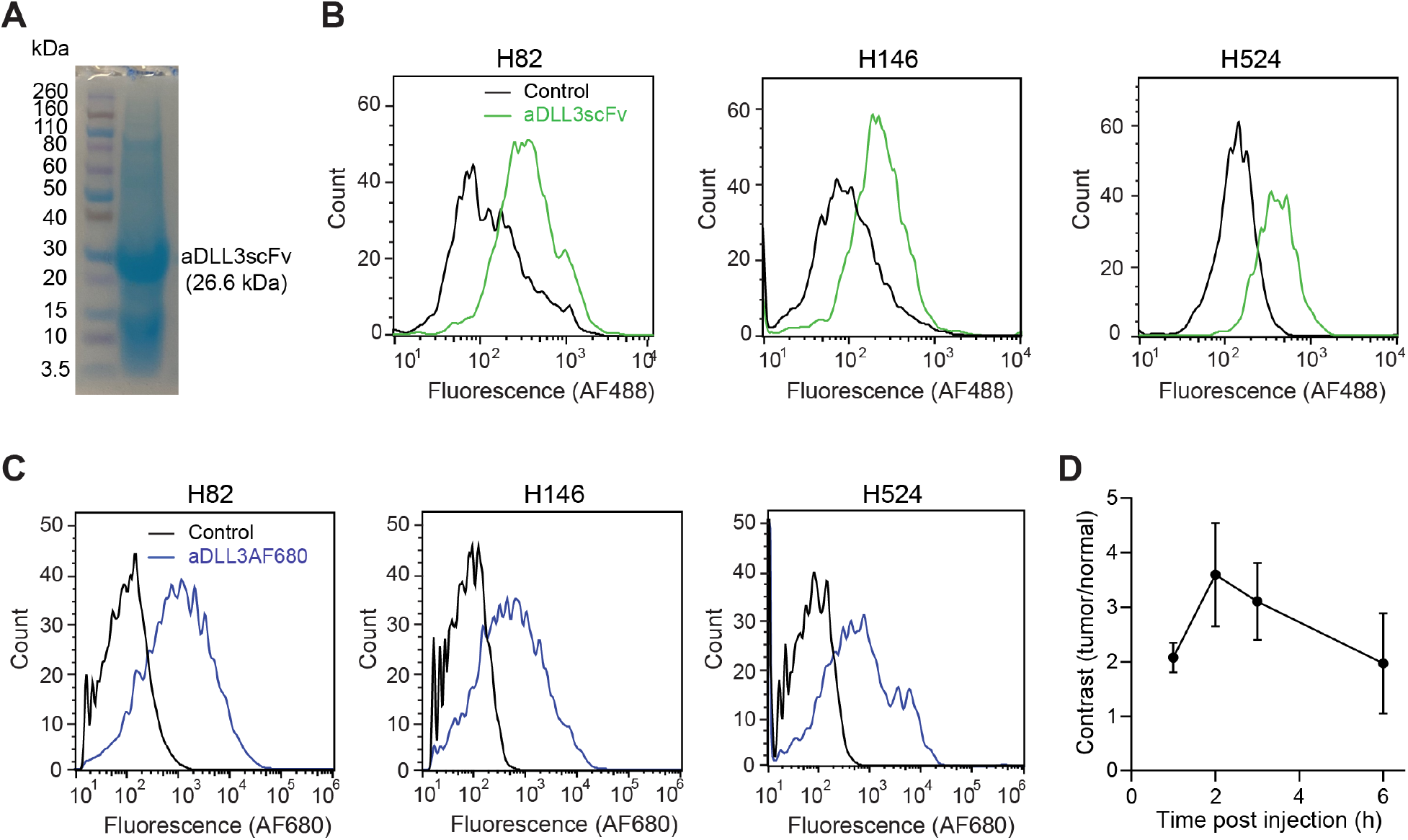
Anti-DLL3 scFv binds to human SCLC cells *in vitro* and *in vivo*. (**A**) SDS-PAGE gel stained with SimplyBlue Safe Stain showing aDLL3scFv purified using nickel-NTA affinity column. (**B**) Representative flow cytometry histograms showing binding of human SCLC cells to His-tagged aDLL3scFv, which was detected using rabbit anti-His antibody and AF488 labeled anti-rabbit antibody. (**C**) Representative flow cytometry histograms showing binding of aDLL3AF680 to human SCLC cells. (**D**) Quantification of imaging contrast reported as the ratio of fluorescent signals for tumor versus normal tissue. Error bars represent s.d., n = 3.

**Figure S3 related to Figure 3.**
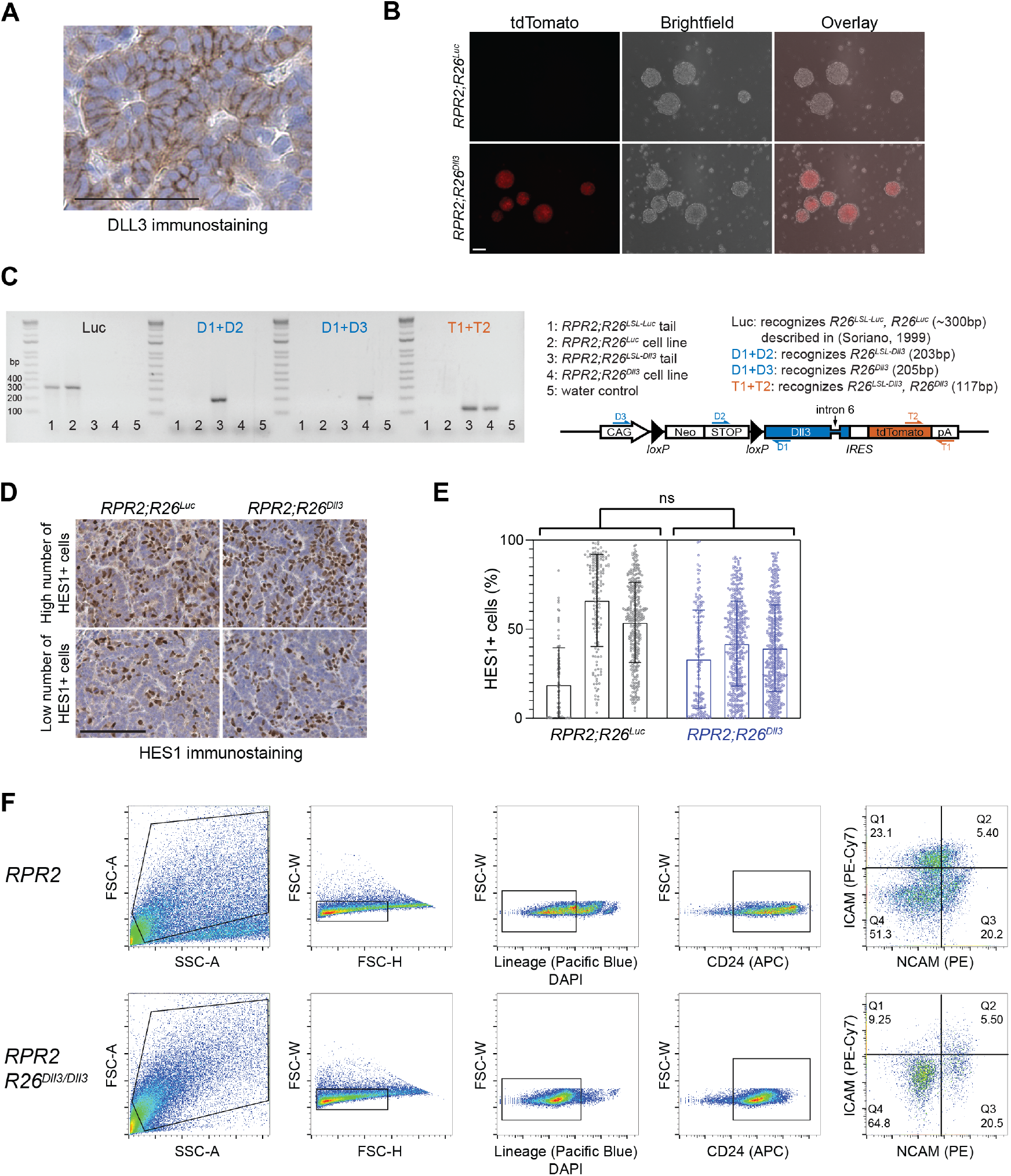
Characterization of a new mouse allele with inducible expression of DLL3. (**A**) Representative DLL3 immunostaining in an *RPR2;R26^Luc^* mutant lung tumor section. Scale bar 50μm. (**B**) Representative tdTomato fluorescence and brightfield images of cell lines derived from *RPR2;R26^Luc^* (top) or *RPR2;R26^Dll3^* (bottom) mutant lung tumors, scale bar 100μm. (**C**) Genotyping PCR and schematic of allele-specific primer pairs. DNA was extracted from the cell lines shown in (B) and from mouse tails for unrecombined controls. Note that the D1-D3 PCR fragment is too long to be amplified in un-recombined alleles. (**D**) Representative HES1 immunostaining in *RPR2;R26^Luc^* (left) and *RPR2;R26^Dll3^* (right) mutant lung tumor sections. Scale bar 100μm. (**E**) Quantification of (D). Each point is a 100 μm diameter region within a tumor, and each bar represents a section from an individual mouse. p=0.597 by nested t-test. Error bars represent SD. (**F**) Flow cytometry strategy to analyze NCAM^high^ ICAM^low^ and NCAM^low^ ICAM^high^ SCLC cells from RPR2;R26^luc^ (top) and RPR2;R26^Dll3^ (bottom) mutant mouse SCLC tumors.

## Notes

### Summary of Updates

Additional data, figure reformatting, minor main text editing

